# MEG sensor patterns reflect perceptual but not categorical similarity of animate and inanimate objects

**DOI:** 10.1101/238584

**Authors:** Daria Proklova, Daniel Kaiser, Marius V. Peelen

## Abstract

Human high-level visual cortex shows a distinction between animate and inanimate objects, as revealed by fMRI. Recent studies have shown that object animacy can similarly be decoded from MEG sensor patterns. Which object properties drive this decoding? Here, we disentangled the influence of perceptual and categorical object properties by presenting perceptually matched objects (e.g., snake and rope) that were nonetheless easily recognizable as being animate or inanimate. In a series of behavioral experiments, three aspects of perceptual dissimilarity of these objects were quantified: overall dissimilarity, outline dissimilarity, and texture dissimilarity. Neural dissimilarity of MEG sensor patterns was modeled using regression analysis, in which perceptual dissimilarity (from the behavioral experiments) and categorical dissimilarity served as predictors of neural dissimilarity. We found that perceptual dissimilarity was strongly reflected in MEG sensor patterns from 80ms after stimulus onset, with separable contributions of outline and texture dissimilarity. Surprisingly, when controlling for perceptual dissimilarity, MEG patterns did not carry information about object category (animate vs inanimate) at any time point. Nearly identical results were found in a second MEG experiment that required basic-level object recognition. These results suggest that MEG sensor patterns do not capture object animacy independently of perceptual differences between animate and inanimate objects. This is in contrast to results observed in fMRI using the same stimuli, task, and analysis approach: fMRI showed a highly reliable categorical distinction in visual cortex even when controlling for perceptual dissimilarity. Results thus point to a discrepancy in the information contained in multivariate fMRI and MEG patterns.

## Introduction

Since their successful application in fMRI research, multivariate analysis methods have recently been applied to MEG and EEG data to gain insight into the temporal dynamics of visual and cognitive processing. A replicable finding in this rapidly growing literature is the finding that MEG sensor patterns carry information about object category (e.g., animacy), peaking around 150-250 ms after stimulus onset (for review, see Contini et al., 2017). Similar category distinctions have been observed in high-level visual cortex using fMRI (Grill-Spector and Weiner, 2014). What are the object properties that drive this decoding, and are these the same in fMRI and MEG?

Animate objects differ from inanimate objects in terms of their characteristic shapes and other category-associated visual features. These feature differences are reflected in behavioral measures of perceptual similarity, such that within- and between-category perceptual similarity can be used to accurately predict the time it takes observers to categorize an object as animate or inanimate (Mohan and Arun, 2012). These perceptual differences likely contribute to categorical distinctions in MEG, considering that object shape and perceptual similarity are strongly reflected in MEG and EEG patterns (Isik et al., 2014; Coggan et al., 2016; Wardle et al., 2016). Furthermore, MEG animacy decoding strength is closely related, at the exemplar level, to categorization reaction time (Ritchie et al., 2015), likely reflecting the exemplar’s perceptual typicality of the category it belongs to (Mohan and Arun, 2012). Together, these studies raise the possibility that category information in MEG patterns primarily reflects perceptual differences between animate and inanimate objects.

However, animate and inanimate objects also differ in other aspects. For example, animals are agents capable of moving by themselves, a property that we rapidly associate with animals even when these are viewed as static pictures. Additionally, we perceive (most) animals as entities with goals, intentions, beliefs, and desires. Animate and inanimate objects also invite different actions on the part of the observer. For example, unlike animals, inanimate objects are often things with specific functions and manipulation patterns, such as tools, musical instruments, or clothing. Animate and inanimate objects are thus associated with different actions, functions, and other higher-order properties. fMRI studies have provided evidence that category selectivity in visual cortex, for example for tools, partly reflects these higher-order associations (for reviews, see Amedi et al., 2017; Peelen and Downing, 2017). This raises the possibility that category information in MEG patterns similarly reflects non-perceptual differences between animate and inanimate objects.

One way to dissociate between these accounts is to compare neural responses to animate and inanimate objects that are perceptually matched in terms of their 2D shape profile (e.g., snake and rope). Recent fMRI studies adopting this approach revealed that category-specific responses in parts of ventral temporal cortex (VTC) are not fully reducible to perceptual differences (Macdonald and Culham, 2015; Bracci and Op de Beeck, 2016; Bryan et al., 2016; Proklova et al., 2016). In a previous fMRI study (Proklova et al., 2016), we found that activity in large parts of the visual cortex reflected the perceptual similarity of the objects, independent of object category. But importantly, the animacy distinction in parts of VTC was preserved for objects that were closely matched in terms of perceptual similarity. If MEG activity in the 150-250 ms time window corresponds to category-selective fMRI activity in these VTC regions, animacy information should be preserved in MEG as well.

In contrast to this prediction, we report here that the matching for perceptual similarity of objects abolished animacy information in MEG: while representational similarity in MEG sensor patterns strongly reflected the perceptual similarity between the objects, as well as similarity in their outline and texture properties, it did not reflect the categorical distinction between animate and inanimate objects at any time point.

## Materials and Methods

### Participants

Twenty-nine participants with normal or corrected-to-normal vision took part in one of two experiments, including 14 volunteers who participated in Experiment 1 (5 females, mean age = 25.6 years, SD = 4 years) and 15 volunteers who participated in Experiment 2 (8 females, mean age = 25.2 years, SD = 3.1 years). All participants provided informed consent and received monetary compensation for their participation. The experimental protocols were approved by the Ethical Committee of the University of Trento, Italy.

### Stimuli

The stimulus set in both experiments was identical to the one used in an earlier fMRI study (Proklova et al., 2016) and consisted of 16 objects (8 animate and 8 inanimate) divided into 4 shape sets (Figure 1A). Each shape set consisted of 2 animals and 2 inanimate objects that were matched for overall shape features (e.g. snake-rope). In addition, four exemplars of each stimulus were used, resulting in a total of 64 stimuli (see Figure 1C for the full stimulus set). In all analyses, we averaged across the 4 exemplars of each stimulus conditions (i.e., there were 16 conditions in total). In Experiment 1, one additional visual stimulus (a hammer, see Figure IB) was used to serve as an oddball target. All stimuli were matched for luminance and contrast using the SHINE toolbox (Willenbockel et al., 2010).

**Figure 1.**
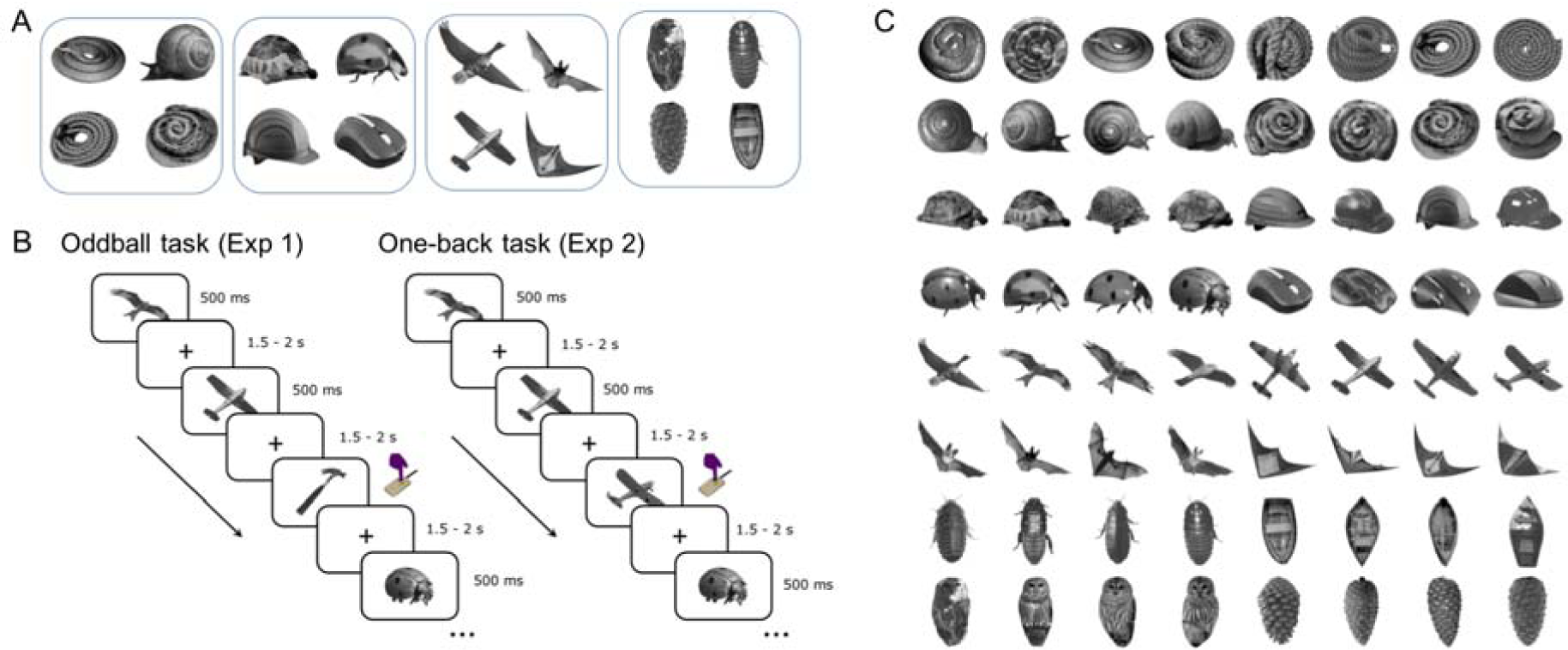
Stimuli and MEG paradigms. (A) Example stimuli of the 16 conditions, grouped in shape clusters. (B) In Experiment 1, individual stimuli were presented centrally for 500 ms, followed by a variable ISI of 1.5 -2 s. Participants were asked to press the response button and blink whenever they sow a hammer. In Experiment 2, the procedure was largely identical to Experiment 1, except that participants performed a one-back task, pressing the button when two images of the same object type (e.g., two different planes) appeared on two consecutive trials. (C) Full stimulus set used in the experiments.

### Experiment 1 and 2: Procedure

Participants viewed the visual stimuli while sitting in the dimly lit magnetically shielded room. The stimuli were projected on a translucent screen located 150 cm from the participant. The stimuli were presented centrally on the uniformly gray background and approximately spanned a visual angle of 8 degrees. Stimulus presentation was controlled using the Psychophysics Toolbox (Brainard, 1997). Participants completed 12 experimental runs, with each of the 64 stimuli appearing exactly twice in each run, in random order. The stimuli were presented for 500 ms, followed by a variable inter-stimulus interval ranging from 1.5 s to 2 s.

In Experiment 1, participants were instructed to maintain fixation and to press the response button and blink each time they saw a picture of a hammer. This target appeared on 20% of trials (32 target trials, randomly distributed over the run), resulting in 160 trials in total per run. Target trials were not analyzed. In Experiment 2, participants were asked to perform a one-back task, pressing the response button whenever an image of the same object type appeared on two consecutive trials (e.g. two different snakes). Twelve repetition trials were inserted at random points within each run, resulting in 140 trials per run. Repetition trials were not analyzed.

### MEG acquisition and preprocessing

Electrophysiological recordings were obtained using an Elekta Neuromag 306 MEG system (Elekta Neuromag systems, Helsinki, Finland) equipped with 204 gradiometers and 102 magnetometers. Signals were sampled continuously at 1000 Hz rate and band-pass filtered online between 0.1 and 330 Hz. The data were then preprocessed offline using MATLAB and the Fieldrip Toolbox (Oostenveld et al., 2011). Data from all runs were concatenated and trials were epoched from 100 ms before to 500 ms after stimulus onset (for one follow-up analysis, trials were epoched from 100 ms before to 800 ms after stimulus onset). Trials containing eye-blinks and other movement-related artifacts were discarded from further analysis based on visual inspection. The signal was baseline-corrected with respect to the pre-stimulus window and downsampled to 100 Hz to reduce the processing time and increase the signal-to-noise ratio (Grootswagers et al., 2017).

### Pairwise decoding analysis

For each time point, we trained linear discriminant analysis (LDA) classifiers on response patterns across sensors to discriminate between all possible pairs of stimuli. For the classification analysis, trials were randomly assigned to four independent chunks, with an equal number of trials for each condition per chunk. Classifiers were trained on data from three chunks and tested on the fourth chunk; this analysis was repeated four times, with each chunk serving as the test set once. To reduce trial-to-trial noise and increase the reliability of data supplied to the classifier, new, “synthetic” trials were created by averaging individual trials (Grootswagers et al., 2017): for every condition and chunk we randomly picked 25% of the original trials and averaged the data across them. This procedure was repeated 500 times (with the constraint that no trial was used more than one time more often than any other trial), producing 500 new “synthetic” trials (with each one being an average of up to 12 trials, depending on trial numbers left after preprocessing) for each condition and chunk that were then supplied to the classifier. The resulting decoding accuracy time course was smoothed with an averaging filter spanning 3 time points (i.e., 30 ms).

The pair-wise decoding analysis yielded a time course of classification accuracies for each pair of stimuli, reflecting the neural dissimilarity of each pair at every time point.

### Representational similarity analysis

In order to investigate the independent contributions of perceptual properties and object category to the MEG signal, we used representational similarity analysis (RSA; Kriegeskorte et al., 2008). Following the analysis approach used in a previous fMRI study (Proklova et al., 2016), the neural dissimilarity (measured with MEG) at each time point was modeled as a linear combination of category and perceptual dissimilarity. The procedures used to obtain the neural dissimilarity matrices as well as category and perceptual dissimilarity matrices are described below.

### Category dissimilarity

A category dissimilarity matrix was constructed by assigning zeroes (minimum dissimilarity) to elements of the matrix that corresponded to pairs of objects belonging to the same category and ones (maximum dissimilarity) for pairs of objects from different categories.

### Perceptual dissimilarity

To be able to measure the contribution of perceptual dissimilarity to the MEG signal, we created predictor matrices that reflected the perceived similarity of the objects for a set of independent observers. In separate behavioral visual search experiments (see Proklova et al., 2016 for a detailed reporting of these experiments), three perceptual dissimilarity matrices were obtained. In one experiment, overall perceptual dissimilarity was quantified using the stimuli used in the MEG experiment. In two other experiments, we sought to dissociate the influence of outline and texture properties of the stimuli by using outline silhouettes and texture patches as stimuli (Figure 2). In all three experiments, participants detected an oddball target among an array of identical distractors (Figure 2) by indicating whether the target was on the right or on the left of the midline. In this task, the identity or category of the target is irrelevant; the task of the participants is simply to indicate where the different-looking stimulus is located (Mohan and Arun, 2012). This was done for all possible target-distractor pairs among the stimuli. For each pair of stimuli, the corresponding entry in the perceptual dissimilarity matrix is given as the inverse reaction time (1/RT) in the visual search task, using one of the stimuli as a target and another as a distractor (Figure 2). For perceptually similar target and distractor pairs, the reaction times will be slower (such as when searching for a snake among ropes), while for perceptually dissimilar target and distractor pairs the reaction times will be faster (such as when looking for a snake among planes).

**Figure 2.**
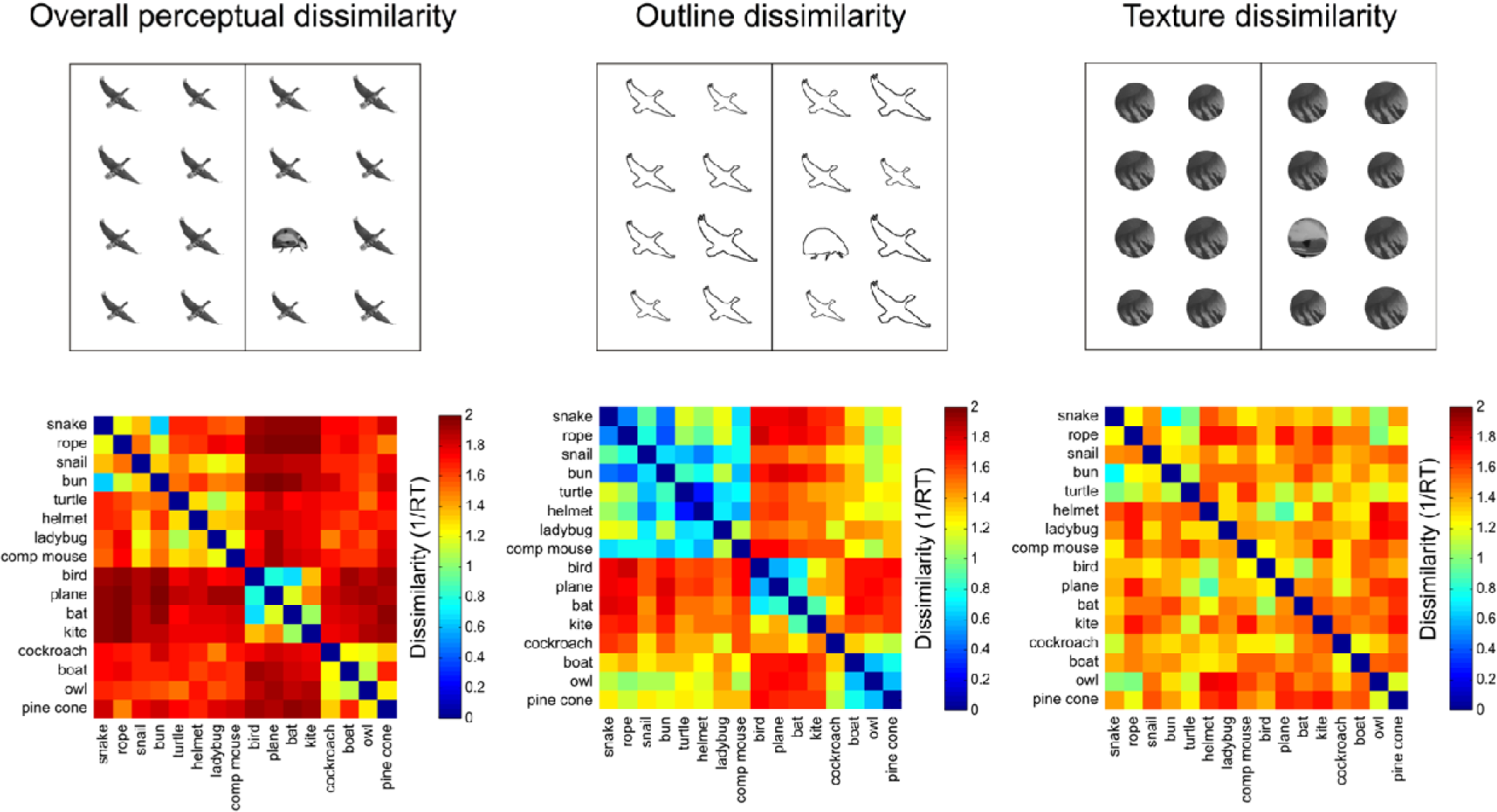
Perceptual dissimilarity measures. Three behavioral visual search experiments were used to measure different aspects of pairwise perceptual dissimilarity of the stimuli. In each experiment, participants had to locate the target among an array of identical distractors. The inverse reaction time in this task is taken as a measure of perceptual dissimilarity between the target and distractor. Dissimilarity matrices are shown for the three behavioral experiments, guantifying overall perceptual dissimilarity (left panel), outline dissimilarity (center), and texture dissimilarity (right panel).

### Relationship between the predictor matrices

Prior to the RSA, we assessed dependencies among the different predictor matrices. Correlating the perceptual dissimilarity matrices with the category dissimilarity matrix revealed that category dissimilarity was not related to the other three predictors (r=-0.08, r=-0.07, and r=-0.06 for overall, outline, and texture dissimilarity, respectively), indicating that the matching of visual properties between animate and inanimate objects was successful. As expected, the three perceptual dissimilarity predictors were related to each other: Overall perceptual dissimilarity correlated with both outline dissimilarity (r=0.82) and texture dissimilarity (r=0.32). Outline and texture dissimilarity were comparably less correlated (r=0.18), suggesting that they capture different aspects of overall perceptual dissimilarity. Interestingly, a linear combination of outline dissimilarity (75% weighting) and texture dissimilarity (25% weighting) closely resembled the overall perceptual dissimilarity (see Proklova et al., 2016, for details). Considering this interdependency between the perceptual dissimilarity matrices, we performed two separate regression analyses: one with category and overall perceptual dissimilarity as predictors, and one with category dissimilarity, outline dissimilarity and texture dissimilarity as predictors.

### Neural dissimilarity

Following the approach used in previous MEG studies, we used pairwise decoding accuracy as a measure of neural dissimilarity, where higher decoding accuracy corresponds to greater neural dissimilarity (Cichy et al., 2014; Wardle et al., 2016). Classifier details are described in the Pairwise decoding analyses section. The classifier accuracy at each time point was assessed as the percentage of correctly classified trials (with chance performance being 50%) and was used as a measure of neural dissimilarity between the two conditions. This classification procedure was performed for all pairs of conditions, resulting in a 16 × 16 neural dissimilarity matrix for each time point.

In addition, as an alternative neural dissimilarity measure we used linear discriminant t-value (LD-t), a version of cross-validated Mahalanobis distance (Nili et al., 2014; Walther et al., 2016). To obtain an LD-t value for each pair of stimuli, the data were divided into two parts (training and testing set), and an LDA classifier was trained to discriminate between the two stimuli (see above). The testing set was then projected on the discriminant dimension, and the t-value was computed describing how well the two stimuli are discriminated. This measure was calculated using the RSA toolbox (Nili et al., 2014). The 16 × 16 LD-t neural dissimilarity matrix was obtained for each time point.

Neural dissimilarity was computed for magnetometers and gradiometers separately, and all subsequent analyses are reported for both sensor types. Previous MEG studies on visual category decoding have either used both sensor types together (e.g., Cichy et al., 2014), only magnetometers (e.g., Kaiser et al., 2016a; 2016b), or only gradiometers (e.g., Ritchie et al., 2015). Here, we decided to report data from magnetometers and gradiometers separately to demonstrate consistency of the results across sensor types (similar results were obtained when analyzing all sensors together).

### Modelling the neural dissimilarity

After constructing neural dissimilarity matrices for each time point, as well as category and perceptual dissimilarity matrices, we performed two regression-based representational similarity analyses (RSA) to determine the contribution of category dissimilarity and different aspects of perceptual dissimilarity to the MEG signal. All dissimilarity matrices were z-scored before estimating the regression coefficients. In the first analysis, the neural dissimilarity at each time point was modeled as a linear combination of category and overall perceptual dissimilarity (Figure 3A). This analysis was performed separately for each participant, resulting in two regression weights for each time point per participant. In the second analysis, the neural dissimilarity at each time point was modeled as a linear combination of category, outline, and texture dissimilarity (Figure 3B). This produced three beta weights per participant for each time point. The time courses of the resulting beta estimates for each predictor were then tested against zero.

**Figure 3.**
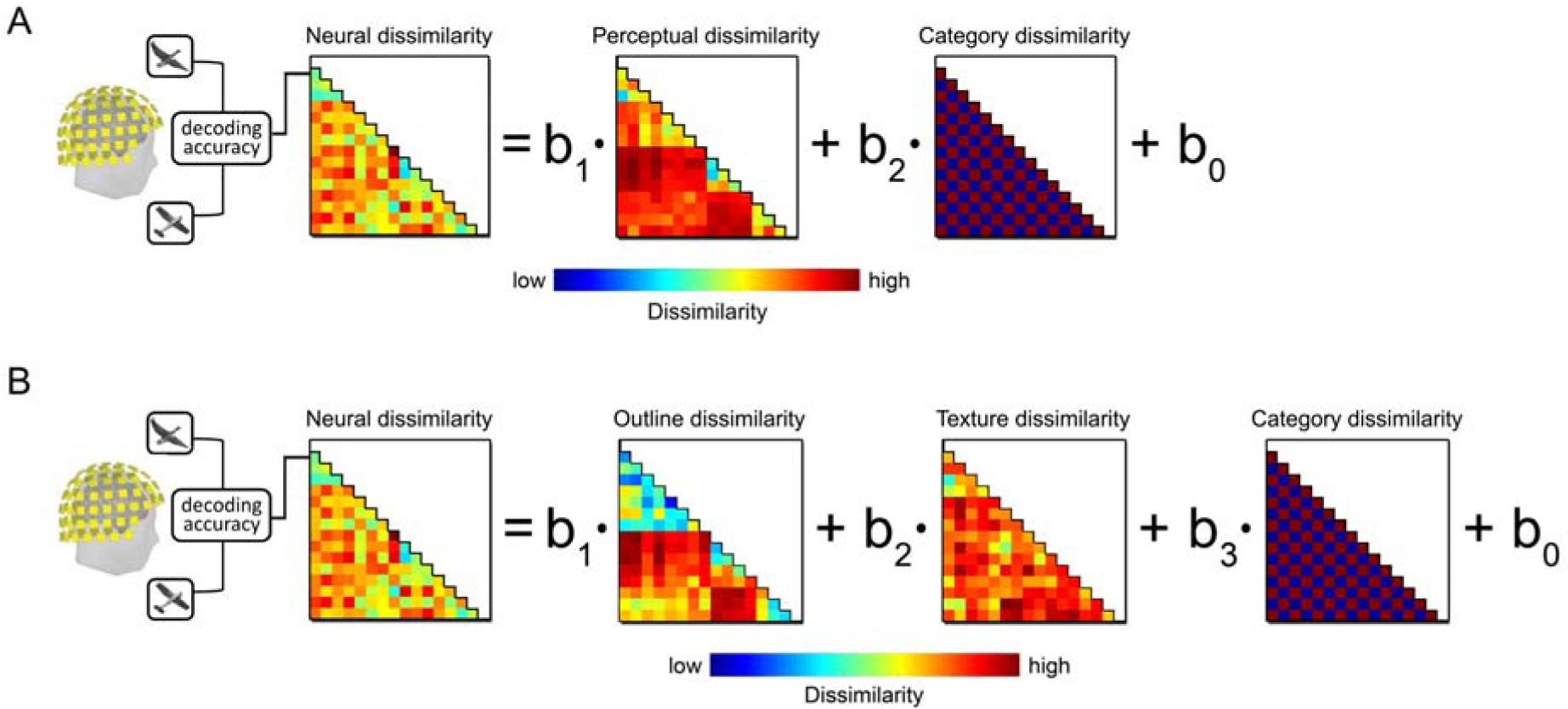
Representational similarity analysis. (A) For the first RSA, at each time point, the neural dissimilarity matrix was constructed by calculating the pairwise decoding accuracy for all pairs of stimuli. This neural dissimilarity matrix was then modeled as a linear combination of overall perceptual dissimilarity and category dissimilarity. (B) For the second RSA, the neural dissimilarity at each time point was modeled using outline, texture and category dissimilarity as predictors.

### Sensor-space searchlight analysis

To identify time periods and sensors in which the contribution of category or perceptual dissimilarity to neural dissimilarity was significant, we performed a sensor-space searchlight analysis. Using the same approach as in the previous analyses, we ran two representational similarity searchlights. For these analyses, the pairwise decoding analysis was repeated for local sensor neighborhoods often sensors each. For each sensor location, neighborhoods were defined by selecting the sensor and its nine nearest neighboring sensors in the MEG gradiometer configuration; individual neighborhoods were thus overlapping in sensor space. For each sensor neighborhood the neural dissimilarity was computed for every post-stimulus time point. Then, to reduce the number of statistical comparisons, the neural dissimilarity matrices for each neighborhood were averaged in time bins of 50 ms, ranging from stimulus onset to 500 ms poststimulus (resulting in 10 time bins).

The resulting neural dissimilarity matrices were then modeled at each time window as (1) a linear combination of category dissimilarity and overall perceptual dissimilarity, resulting in two beta estimate maps, and (2) a linear combination of category dissimilarity, outline dissimilarity, and texture dissimilarity, producing three beta estimate maps. The resulting beta estimates were mapped onto a scalp representation. The scalp maps for each predictor were then averaged across participants and tested against zero.

### Statistical analysis

For all tests, statistical significance was assessed using the threshold-free cluster enhancement procedure (TFCE) (Smith and Nichols, 2009) with default parameters, using multiple-comparison correction based on a sign-permutation test (with null distributions created from 10,000 bootstrapping iterations) as implemented in the CoSMoMVPA toolbox (Oosterhof et al., 2016). The threshold was set at Z > 1.64 (i.e., p < 0.05, one-tailed), as further clarified in the individual sections. Significance in the searchlight analysis was assessed separately for each time window using the TFCE procedure to reveal the sensors in which the contribution of a particular predictor to the neural dissimilarity was significantly above chance.

## Results

### Behavioral results

Participants were very accurate in the oddball detection task of Experiment 1 (mean = 97%, SD = 2%) and the one-back repetition detection task of Experiment 2 (mean = 94%, SD = 1%). Responses were faster in Experiment 1 (mean = 0.47 s, SD = 0.06 s) than Experiment 2 (mean = 0.73 s, SD = 0.12 s).

### Decoding stimulus conditions

To assess the quality of the MEG data and evaluate whether stimulus condition could be decoded from the MEG sensor patterns, we averaged the off-diagonal elements of the MEG dissimilarity matrix for each time point. This was done separately for Experiments 1 and 2, as well as for each of the two sensor types (magnetometers and gradiometers), producing four time courses of average pairwise stimulus decodability (Figure 4). In both experiments, and for both sensor types, decoding was at chance until about 50 ms, after which the decoding curve rose sharply, becoming significant at 60 ms (or at 70 ms for magnetometers in Experiment 2) and peaking at 120 ms. This pattern is similar to the pattern observed in previous studies (for review, see Contini et al., 2017). For both sensor types, average pairwise decoding accuracy was higher in Experiment 2, suggesting that the demands of a one-back task (e.g., enhanced attention to the objects, deeper object processing, and involvement of working memory) increases neural discriminability. Overall, these results replicate earlier MEG decoding studies and show that the individual stimuli could be decoded successfully from the MEG signal.

**Figure 4.**
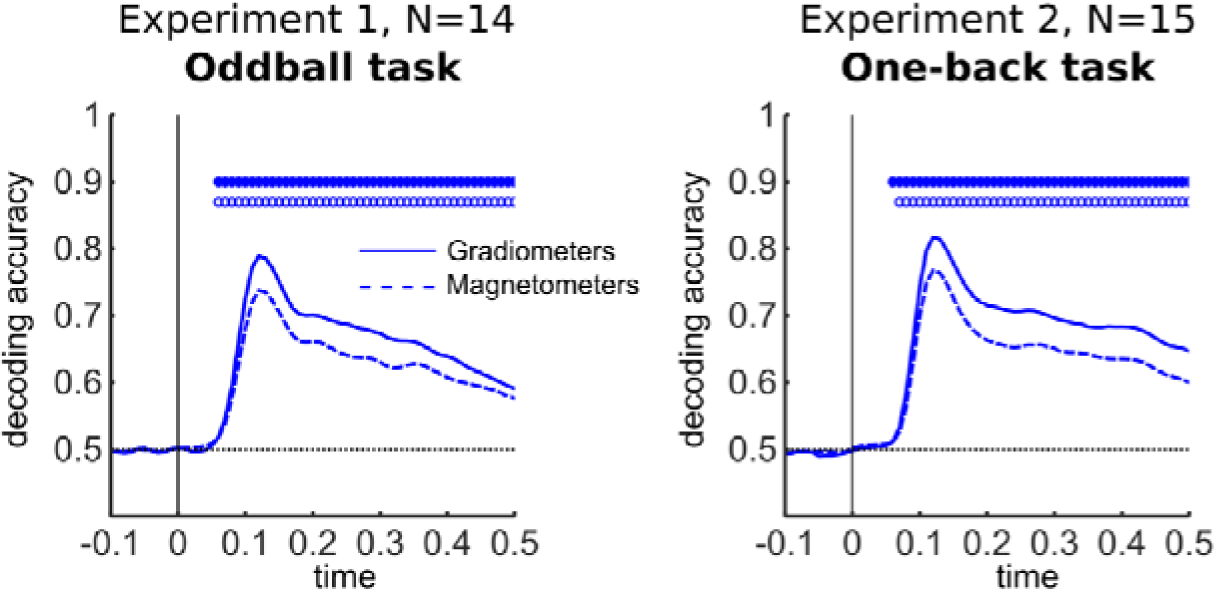
Time courses of average decodability of all stimulus pairs. Average pairwise stimulus decodability in Experiment 1 (left panel) and Experiment 2 (right panel), shown separately for two sensor types (solid line for gradiometers and dashed line for magnetometers). Circles indicate time bins where decoding accuracy was significantly greater than chance (0.5), with filled circles showing significant time points for gradiometers and empty circles for magnetometers.

### Representational similarity analysis

To examine the separate contribution of overall perceptual dissimilarity (based on RTs in the visual search task) and category dissimilarity to neural dissimilarity, we performed a representational similarity analysis (RSA) modelling the neural dissimilarity at each time point as the linear combination of these two predictors (see Materials and Methods; Figure 3A). This analysis produced one beta estimate time course for each predictor. The results of this analysis for both experiments are shown in Figure 5A, separately for the two types of sensors. For both magnetometers and gradiometers, the beta estimate for overall perceptual dissimilarity (shown in red) reached significance at 80 ms, peaking at 130 ms. The beta estimate for category dissimilarity (shown in blue) did not reach significance at any time point. In a second analysis, the neural dissimilarity was modeled as a combination of category, outline, and texture dissimilarity (Figure 3B). This analysis produced three time courses that are shown in Figure 5B. The outline predictor contributed significantly to neural dissimilarity, starting at 80 ms and peaking at 150 ms, followed by a smaller peak at 250 ms. The time course of the texture predictor was significantly above chance starting from 90 ms, with a peak at 130 ms. The beta estimate for category dissimilarity did not reach significance at any time point.

**Figure 5.**
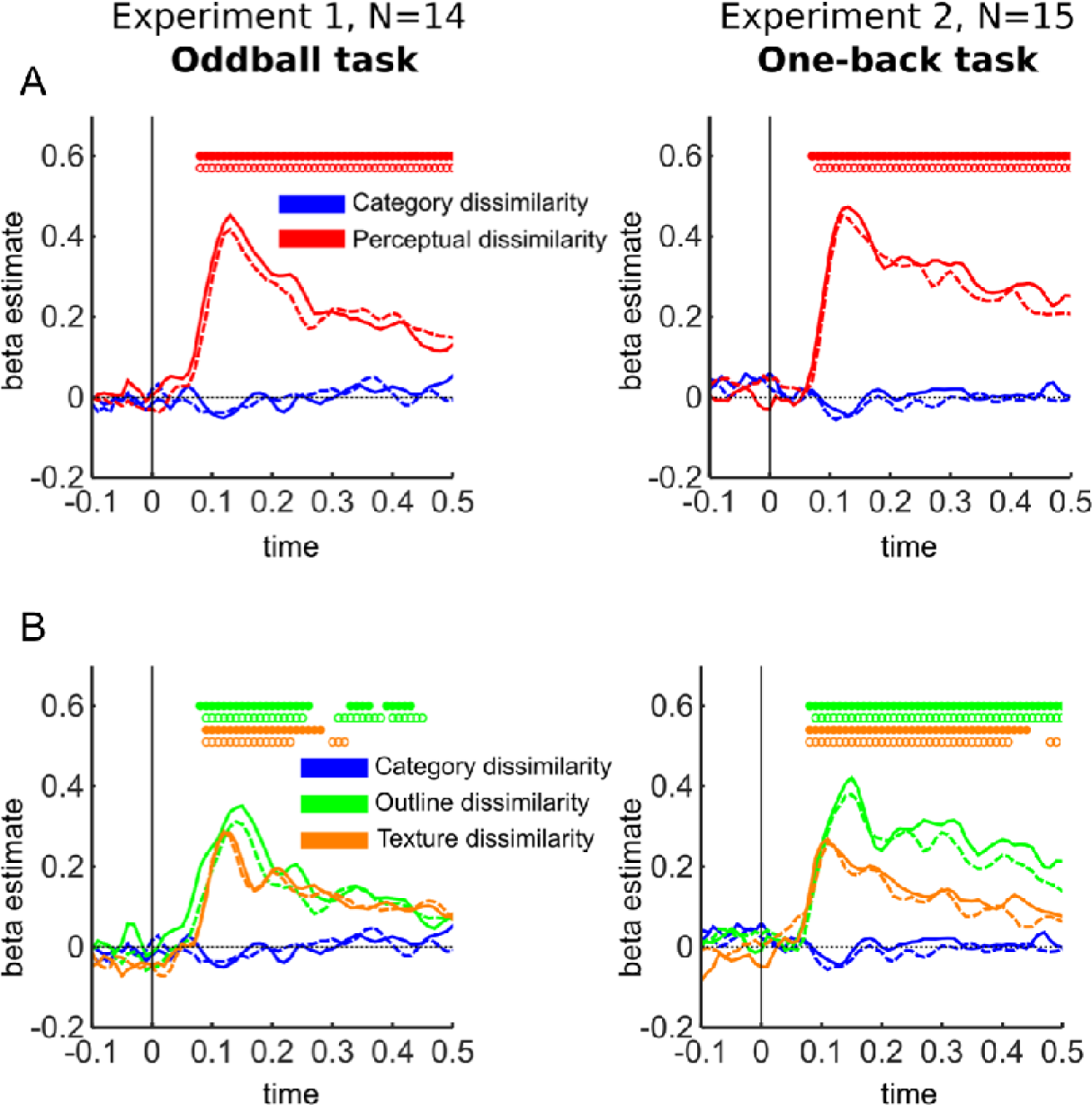
RSA results using decoding accuracy. (A) The time courses of regression weights for two sensor types (solid line for gradiometers and dashed line for magnetometers) showing the contributions of category dissimilarity (in blue) and overall perceptual dissimilarity (in red) to neural dissimilarity for Experiment 1 (left column) and Experiment 2 (right column). (B) The time courses of regression weights reflecting the contributions of category dissimilarity (in blue), outline dissimilarity (in green) and texture dissimilarity (in orange). Circles indicate time bins where beta estimates were significantly greater than zero, with filled circles showing significant time points for gradiometers and empty circles for magnetometers.

Nearly identical results were obtained in Experiment 2, in which subject performed a one-back task (Figure 5, right column). The first RSA again revealed a strong contribution of overall perceptual dissimilarity, peaking at 120 ms for magnetometers and at 130 ms for gradiometers (Figure 5A, right column). In the second RSA, the outline dissimilarity significantly contributed to the neural dissimilarity with a peak at 150 ms for both magnetometers and gradiometers (Figure 5B, right column). Texture (shown in orange) peaked at 110 ms for both types of sensors. In both analyses the category predictor again did not reach significance at any time point, confirming the results of Experiment 1 with a different and more engaging task that required attention to object identity.

It is possible that the previously described measure of neural dissimilarity (i.e., pairwise decoding accuracy) was not sensitive to reveal category effects. fMRI studies have suggested that the cross-validated Mahalanobis distance may be a more reliable measure of neural dissimilarity (Walther et al., 2016). Thus, we repeated all analyses using the neural dissimilarity matrices constructed using this measure (see Materials and Methods). The RSA results using this alternative neural dissimilarity measure are shown in Figure 6. The pattern of results was nearly identical to the one obtained using decoding accuracy.

**Figure 6.**
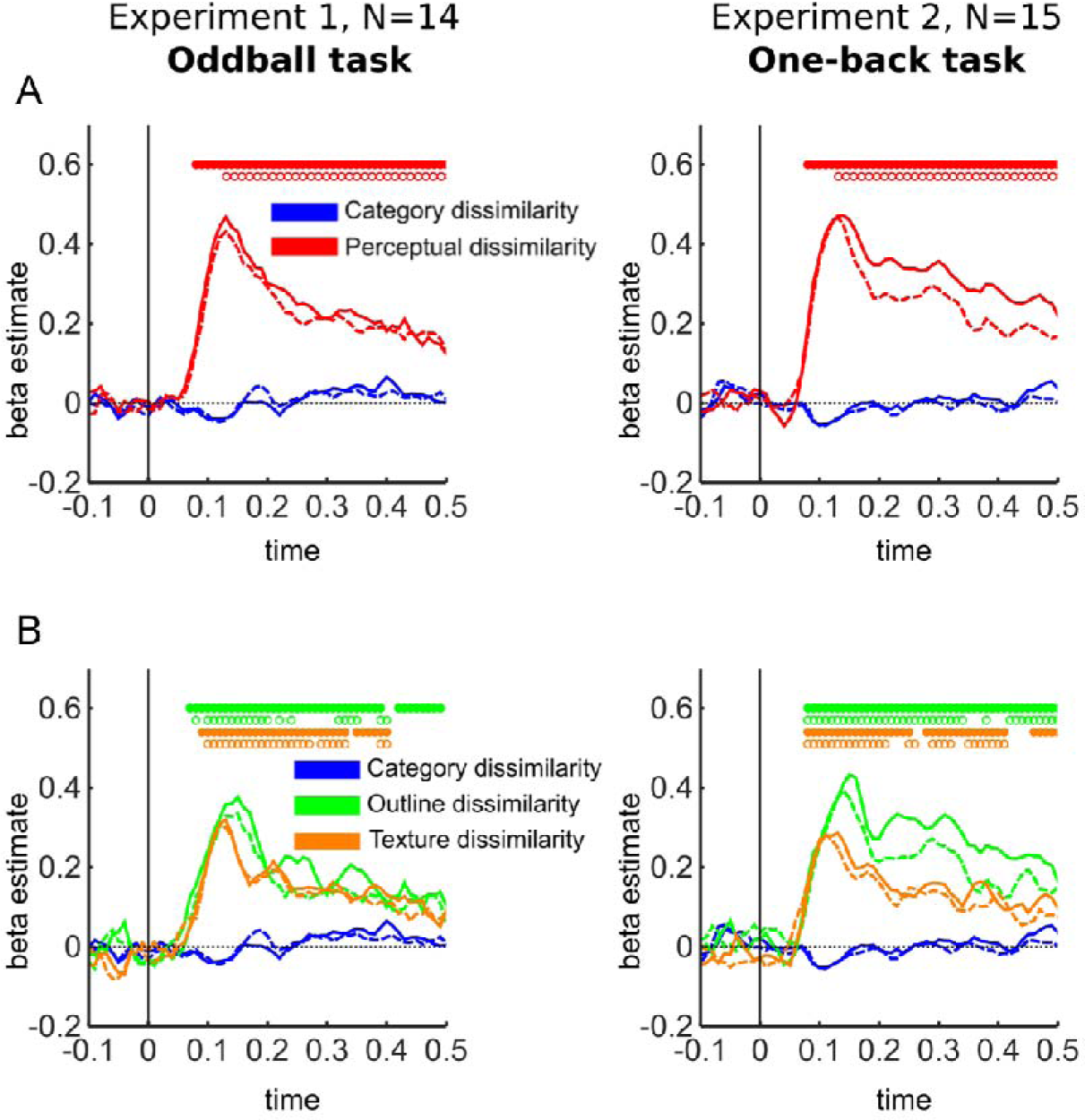
RSA results using cross-validated Mahalanobis distance. (A) Contributions of category dissimilarity (in blue) and perceptual dissimilarity (in red) to neural dissimilarity for two sensor types (solid line for gradiometers and dashed line for magnetometers). Results of Experiment 1 (left panel) and Experiment 2 (right panel). (B) The same analysis for three predictors (outline dissimilarity, texture dissimilarity, category dissimilarity). Circles indicate time bins where beta estimates were significantly greater than zero, with filled circles showing significant time points for gradiometers and empty circles for magnetometers.

Finally, to maximize statistical power to detect category information, we combined the data of the two experiments, resulting in N=29. Moreover, to explore the possibility that category information emerges after 500 ms, we repeated the analysis for a longer time window, until 800 ms after stimulus onset. The results of these analyses are shown in Figure 7. In the first RSA (Figure 7A), the overall perceptual dissimilarity beta estimate peaked at 130 ms for both types of sensors and for both decoding accuracy (left column) and cross-validated Mahalanobis distance (right column) as measures of neural dissimilarity. In the second RSA (Figure 7B), the regression weights for outline dissimilarity peaked at 140 ms for magnetometers and at 150 ms for gradiometers for both dissimilarity measures. Texture dissimilarity beta estimates peaked at 120 ms for both sensor types. Category dissimilarity was not significant at any time point.

**Figure 7.**
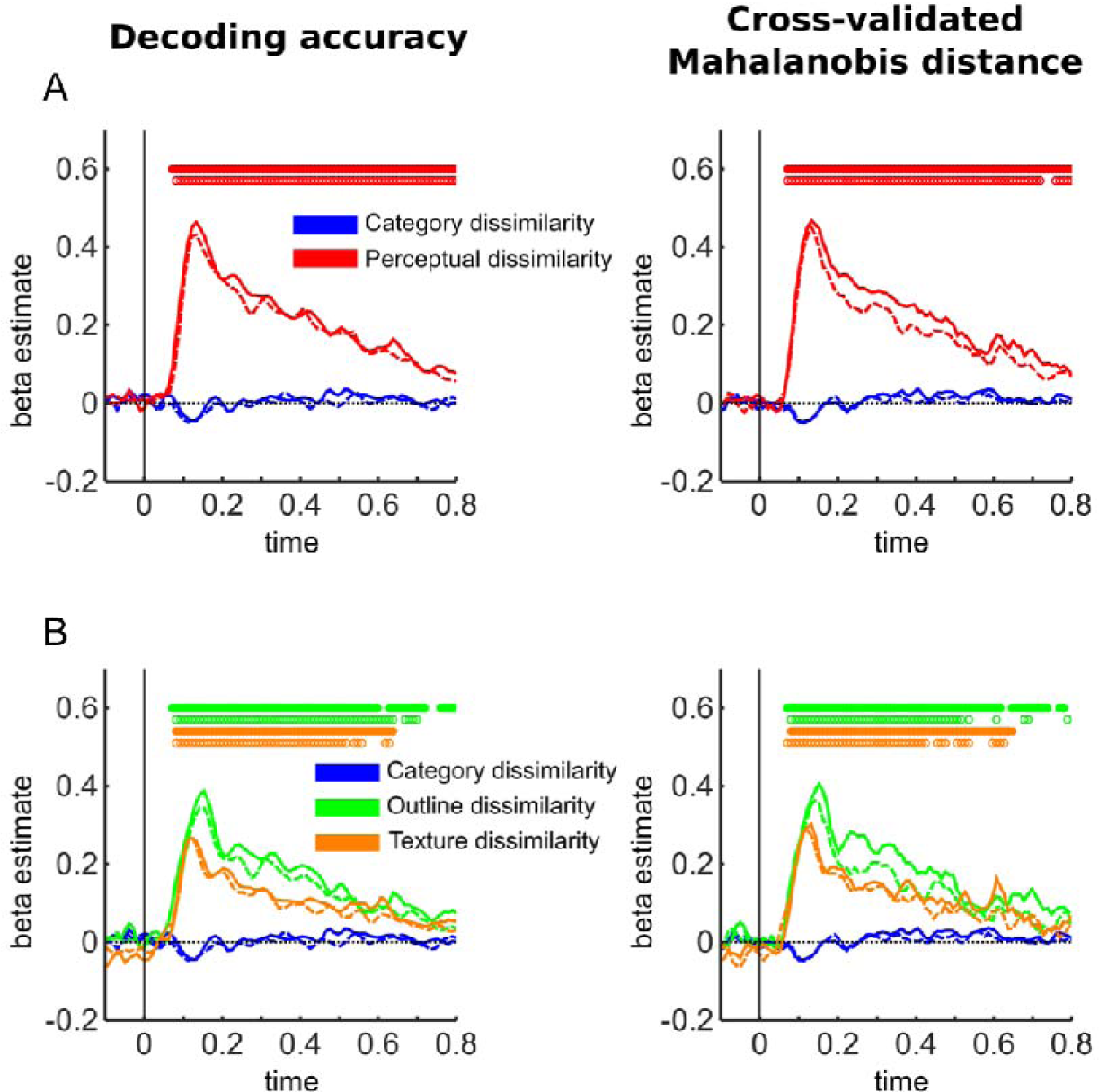
RSA results for both experiments combined (N=29). (A) Contributions of category dissimilarity (in blue) and perceptual dissimilarity (in red) to neural dissimilarity for two sensor types (solid line for gradiometers and dashed line for magnetometers). Results of both experiments combined using decoding accuracy as the measure of neural dissimilarity (left panel), and cross-validated Mahalanobis distance (right panel). (B) The same analysis for three predictors (outline dissimilarity, texture dissimilarity, category dissimilarity). Circles indicate time bins where beta estimates were significantly greater than zero, with filled circles showing significant time points for gradiometers and empty circles for magnetometers.

### Representational similarity searchlight

The absence of category information in the previous analyses could in principle be due to differences in the spatial scales of perceptual and category processing: perhaps perceptual properties are well reflected in coarse patterns across all sensors, whereas more subtle categorical responses are only reflected in localized patterns emerging across a few sensors. We therefore performed a sensor-space representational similarity searchlight analysis. This analysis was done using the data from gradiometers, because these sensors have greater spatial specificity and showed numerically greater pairwise discriminability (e.g., Figure 4). Given that the pattern of results in Experiments 1 and 2 was highly similar we pooled data from Experiments 1 and 2 for the searchlight analysis to maximize power.

For each channel, we defined a neighborhood of ten adjacent channels and computed the neural dissimilarity matrix for this neighborhood (see Materials and Methods). We then used the RSA approach described earlier, modelling the neural dissimilarity for each channel as a combination of category and perceptual dissimilarity. This procedure was repeated for each 50 ms time window (see Materials and Methods), resulting in 10 searchlight maps showing the contribution of perceptual and category dissimilarity to neural dissimilarity at different sensor locations. In accordance with the previous analyses, this analysis did not reveal any sensors or time windows in which the contribution of category dissimilarity to neural dissimilarity was significantly above zero (Figure 8A). In contrast, the contribution of perceptual dissimilarity again reached significance in the 50-100 ms time window and stayed significant throughout all subsequent time windows, starting in a group of posterior sensors and progressing to more anterior sensors at 100-250 ms, peaking at 150-200 ms and gradually receding towards the posterior sensors again (Figure 8B). These results suggest that the MEG neural dissimilarity across multiple channel locations predominantly reflects perceptual, but not category dissimilarity, also when looking at more localized sensor patterns.

**Figure 8.**
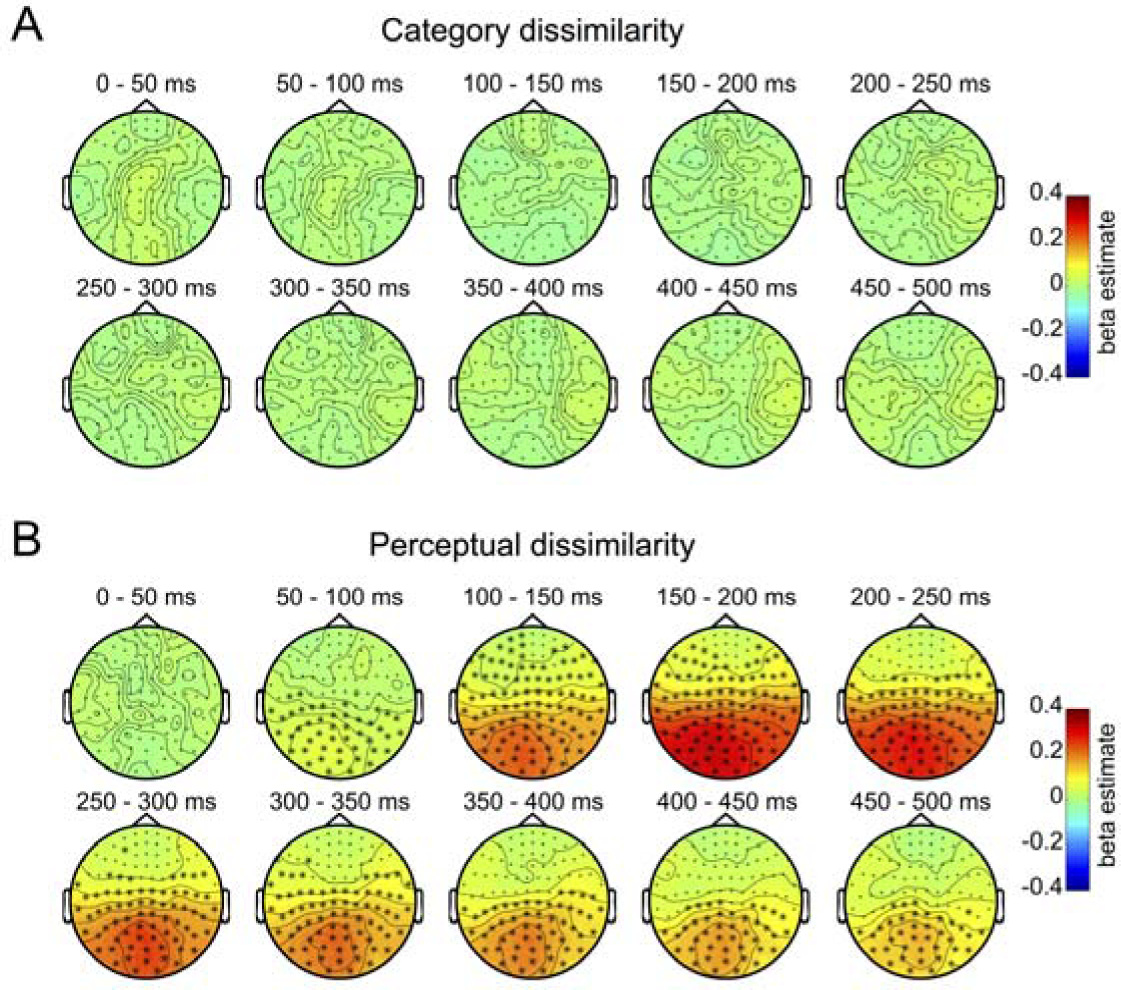
RSA searchlight results: category and perceptual dissimilarity. (A) Searchlight maps showing the regression weights reflecting the contribution of category dissimilarity to MEG patterns across channels (gradiometers) and ten time windows. The beta estimates for category were not significantly above zero in any of the sensors for all time windows. (B) Searchlight maps showing the perceptual dissimilarity regression weights across gradiometers for ten time windows. Asterisks indicate sensor locations where beta estimates were significantly greater than zero.

Finally, we ran a second RSA searchlight, using category, outline and texture dissimilarity as predictors of neural dissimilarity. The results of this analysis are shown in Figure 9. As in the previous analysis, no sensors in any time window exhibited a significant contribution of category dissimilarity to MEG neural dissimilarity (Figure 9A). By contrast, the outline dissimilarity beta estimates reached significance at 50-100 ms after stimulus onset in a group of left posterior sensors, spreading to more anterior sensors at 100-150 ms and peaking at 150-200 ms, staying significant in posterior channels throughout all time windows (Figure 9B). Similarly, texture dissimilarity beta estimates reached significance in a cluster of central posterior sensors at 50-100 ms and stayed significant at multiple channel locations in all the next time windows, peaking at 150-200 ms (Figure 9C).

**Figure 9.**
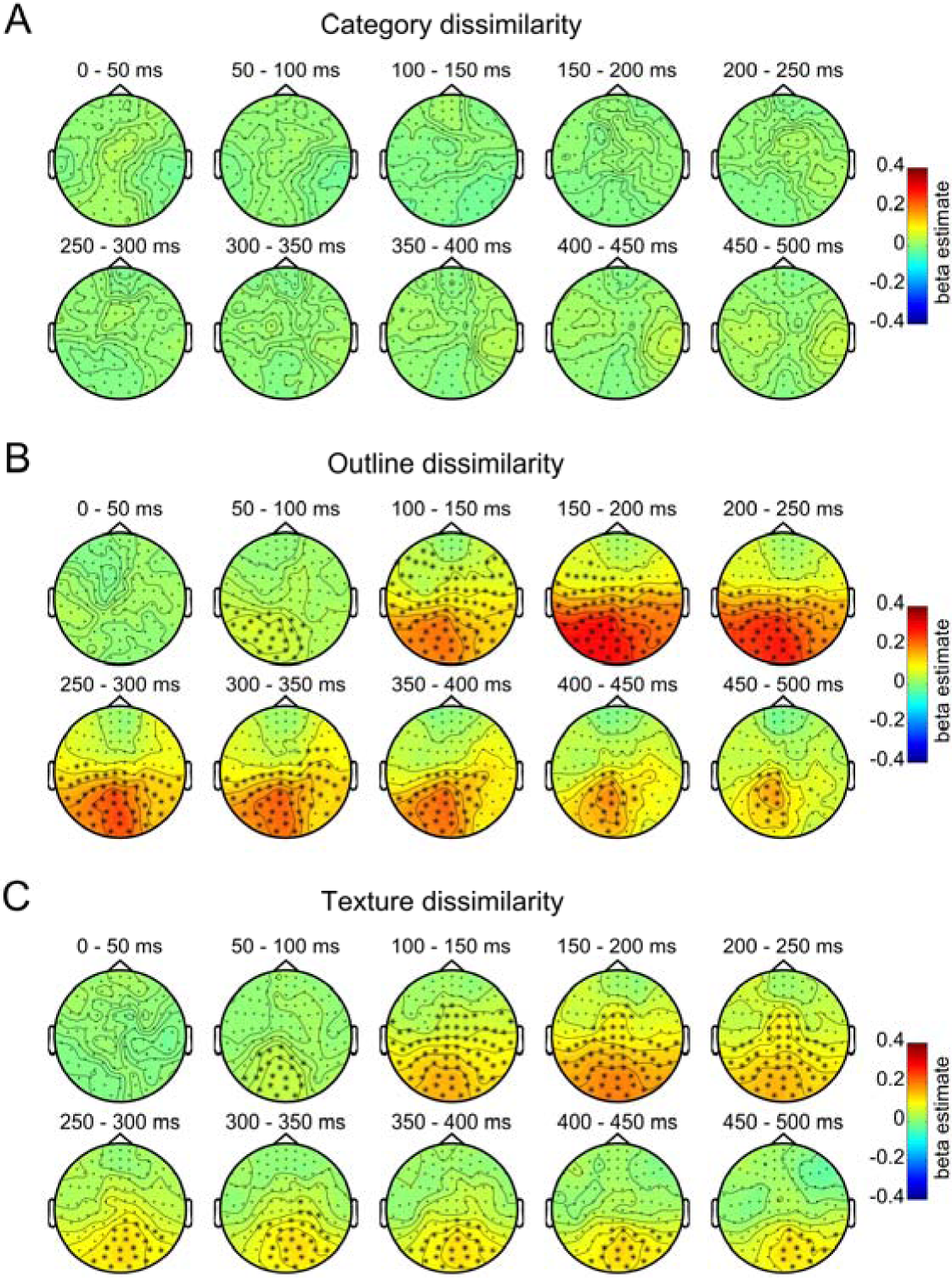
RSA searchlight results: category, outline and texture dissimilarity. (A) Beta estimate maps showing the contribution of category dissimilarity to MEG patterns across channels (gradiometers) and ten 50-ms time windows. The beta estimates for category did not reach significance in any of the sensors or time windows. (B) Beta estimate maps showing the contribution of outline dissimilarity to the MEG signal across gradiometers for ten time windows. (C) Beta estimates reflecting contribution of texture dissimilarity to neural dissimilarity. Asterisks indicate sensor locations where beta estimates were significantly greater than zero.

## Discussion

In this study, we used a carefully designed stimulus set to disentangle categorical from perceptual properties in driving MEG sensor patterns. Replicating previous reports, MEG sensor patterns carried information about individual objects (Carlson et al., 2011; Carlson et al., 2013; van de Nieuwenhuijzen et al., 2013; Cichy et al., 2014; Isik et al., 2014; Clarke et al., 2015; Ritchie et al., 2015; Coggan et al., 2016; Kaiser et al., 2016a). Using representational similarity analysis, we then related MEG neural similarity to the objects’ perceptual and categorical similarity. This analysis revealed the time course of the objects’ overall perceptual similarity as well as the separate contributions of outline shape and inner texture; each of these independently contributed to MEG neural similarity but with overlapping time courses. Contrary to our prediction, there was no category information in the MEG patterns even though participants easily recognized the objects at the basic level, as evidenced by behavioral performance in Experiment 2 and post-experiment debriefing.

These results can be directly compared with those of a recent fMRI study using the same stimulus set, task, and analysis approach (Proklova et al., 2016). Similar to the current MEG findings, outline shape and texture also contributed independently to multi-voxel fMRI patterns in visual cortex, with these regions partly overlapping. However, unlike the current MEG results, fMRI showed a highly reliable contribution of object category to neural similarity in anterior VTC. Our results therefore show that the representation of animacy for shape-matched objects in VTC does not give rise to distinct scalp-level patterns as observed with MEG. It should be noted, however, that considering the fact that the animacy of our stimuli during this task *is* represented in the brain (Proklova et al., 2016), it remains a possibility that other types of analysis (e.g., decoding in source-space; van de Nieuwenhuijzen et al., 2013) or additional preprocessing of the MEG data (e.g., principal component analysis; Grootswagers et al., 2017) could still retrieve this information. Nevertheless, our results suggest that any such information is likely to be very weak relative to information about visual properties.

Our results show that MEG sensor patterns in the 150-250 ms range reflect perceptual properties, including outline shape and inner texture, derived from response times in a visual search task. Previous behavioral work has shown that this perceptual similarity measure can be used to predict animate-inanimate categorization times when using stimuli that naturally confound visual features and category (Mohan and Arun, 2012). For example, a side-view picture of a cow is quickly categorized as animate because it is perceptually relatively similar to other animals (e.g., sheep, horse, dog) and relatively dissimilar from inanimate objects. These findings, combined with our current results, suggest that previous MEG reports of animacy decoding in the 150-250 ms range likely reflected the perceptual differences between animate and inanimate objects. In the current stimulus set, the perceptual similarity measure no longer reflects categorical influences (Proklova et al., 2016), allowing us to dissociate these two components and reveal that MEG sensory patterns at 150-250 ms reflect perceptual object properties rather than object category *perse.*

It should be noted that previous studies provided some evidence for a contribution of semantic object properties to the MEG signal even at relatively early latencies (<250 ms; Clarke et al., 2015; Coggan et al., 2016; Kaiser et al., 2016b). These studies differed from the current study in several important ways. First, none of these studies controlled for perceptual similarity using human judgments or task performance. It is likely that the semantic factors in these studies (e.g., the property “has legs”; Clarke et al., 2015) would still express in perceptual similarity differences in tasks like the visual search task used here. Second, two of these studies included human faces (Coggan et al., 2016) or bodies (Kaiser et al., 2016b). The human visual system is particularly sensitive to such stimuli (Stein et al., 2012), showing highly selective and localized face- and body-selective responses in fMRI (Kanwisher, 2010).

Notably, these face- and body-selective responses are right lateralized in most participants (Willems et al., 2010), thus resulting in distinct scalp topographies (Thierry et al., 2006) that are likely more easily decodable than the bilateral animate-inanimate organization investigated here (no significant lateralization of animacy was observed in Proklova et al., 2016, see their Figure 3B). Similarly, the animate stimuli used in the current study (birds, reptiles, insects) give a lower response than mammals in right-lateralized face- and body-selective regions (Downing et al., 2006), which may reflect their lower score on a proposed animacy continuum (Sha et al., 2015).

These considerations leave open the possibility that MEG sensor patterns contain information about more human-like animals (e.g., mammals), even after shape matching. Future studies should test the decoding of other shape-matched categories to investigate whether lateralization is indeed required for MEG to reveal categorical representations in high-level visual cortex.

To conclude, the current study shows that MEG sensor patterns are highly sensitive to two independent perceptual features (outline shape and texture) of visually presented objects, quantified using behavioral perceptual similarity measures. By contrast, MEG sensor patterns appear insensitive to the conceptual-level distinction between animate and inanimate objects, unlike fMRI voxel patterns in ventral temporal cortex (Proklova et al., 2016). The discrepancy between the information contained in MEG and fMRI patterns may need to be considered when integrating these methods, for example using RSA (Cichy et al., 2016).

## Acknowledgements

The research was supported by the Autonomous Province of Trento, Call “Grandi Progetti 2012”, project “Characterizing and improving brain mechanisms of attention-ATTEND”. This project has received funding from the European Research Council (ERC) under the European Union’s Horizon 2020 research and innovation programme (grant agreement No 725970). D.K. is supported by a DFG grant (KA4683-2/1).

